# Total Chemical Synthesis of Glycosylated TREM2 Ectodomain

**DOI:** 10.1101/2023.04.14.536909

**Authors:** Gayani Wijegunawardena, Erika Castillo, Brandy Henrickson, Regan Davis, Carlo Condello, Haifan Wu

**Affiliations:** Department of Chemistry and Biochemistry, Wichita State University, Wichita, Kansas 67260, United States; Institute for Neurodegenerative Diseases, University of California, San Francisco, CA 94158, United States; Department of Neurology, Weill Institute for Neurosciences, University of California, San Francisco, CA 94158, United States

**Keywords:** TREM2, native chemical ligation, glycosylation, Alzheimer’s disease

## Abstract

Mutations in a microglia-associated gene *TREM2* increase the risk of Alzheimer’s disease. Currently, structural and functional studies of TREM2 mainly rely on recombinant TREM2 proteins expressed from mammalian cells. However, using this method, it is difficult to achieve site-specific labeling. Here we present the total chemical synthesis of the 116 amino-acid TREM2 ectodomain. Rigorous structural analysis ensured correct structural fold after refolding. Treating microglial cells with refolded synthetic TREM2 enhanced microglial phagocytosis, proliferation, and survival. We also prepared TREM2 constructs with defined glycosylation patterns and found that glycosylation at N79 is critical to the thermal stability of TREM2. This method will provide access to TREM2 constructs with site-specific labeling, such as fluorescent labeling, reactive chemical handles, and enrichment handles, to further advance our understanding of TREM2 in Alzheimer’s disease.

## INTRODUCTION

Human genetic studies have identified several genes associated with a higher risk of developing Alzheimer’s disease (AD).^1, 2^ Several of them have been linked to microglia, resident immune cells in the brain, suggesting that microglial dysfunction contributes to the progression of AD.^3-6^ Among these microglial risk genes, *TREM2* (triggering receptor expressed on myeloid cells 2) encodes a cell surface receptor that senses a variety of molecules, including phospholipids, apolipoproteins, oligomeric amyloid β (Aβ), etc.^7-11^ TREM2 signaling is believed to maintain metabolic fitness of microglia and create a neuroprotective barrier in response to AD pathology.^12-14^ However, a partial loss of TREM2 function due to mutations significantly increases AD risk. For example, the R47H mutation increases AD risk by three fold.^15, 16^ Both human genetics and studies of animal models have identified TREM2 as a potential therapeutic target against AD.^13, 17-19^

TREM2 can be proteolytically cleaved by proteases ADAM10/17 between residues H157 and S158 to release a soluble fragment TREM2(19-157) or sTREM2.^20^ Although the exact biological function of sTREM2 is not fully elucidated, multiple lines of evidence support that sTREM2 is a bioactive molecule playing a protective role against Alzheimer’s disease. For example, a recent longitudinal study found that elderly individuals with higher sTREM2 levels in the cerebrospinal fluid (CSF) showed a slower rate of amyloid beta deposition.^21^ In addition, the direct injection of recombinant sTREM2 into the brain of AD transgenic mice or increased sTREM2 expression via adeno-associated viral vectors reduced amyloid pathology and improved memory in AD transgenic mice,^22^ suggesting a therapeutic potential of sTREM2. This sTREM2-mediated protection is dependent on microglia.^22^ We and others have discovered that sTREM2 can inhibit the aggregation of amyloid beta *in vitro* and increase the uptake of aggregated amyloid beta by microglia.^23-25^ Furthermore, sTREM2 was shown to enhance microglia survival and proliferation.^22, 26^

*In vitro* studies of TREM2 to understand the molecular details of its structures and functions require the production of homogeneous TREM2 proteins with site-specific modifications, such as glycosylation, fluorescence labeling, biotinylation, etc. The expression using HEK cells can provide glycosylated TREM2 with highly heterogeneous glycans at residues N20 and N79.^27^ To generate homogeneous TREM2 proteins for structural determination by X-ray crystallography, an N20D mutation has been introduced followed by treatment with endo H to trim glycans at N79 to monosaccharide.^28, 29^ However, the generation of TREM2 constructs with homogeneous glycoforms at both N20 and N79 sites is still not demonstrated. Fluorescence or biotin labeling using reagents targeting lysine residues is a common way to introduce fluorophores or biotin into proteins. However, this process generates a mixture of TREM2 proteins with different levels and sites of labeling. Additionally, lysine residues are critical in TREM2 as they form a basic patch in TREM2 responsible for ligand binding.^28^ Labeling by targeting lysine residues could affect the functions of TREM2. Therefore, an alternative strategy is needed to produce homogeneous TREM2 constructs with site-specific modifications.

Chemical protein synthesis via native chemical ligation connects short to medium-sized peptide fragments into a long polypeptide chain.^30-32^ As each peptide fragment can be individually synthesized via solid-phase peptide synthesis (SPPS), site-specific modifications can be conveniently introduced into proteins. Here, we describe the synthesis of glycosylated TREM2 ectodomain. We applied CD spectroscopy, disulfide mapping, and cross-linking to ensure the right structure fold of synthetic TREM2 ectodomain. Microglia cell-based assays were performed to demonstrate its bioactivity. We further applied this synthetic method to generate well-defined glycoforms of TREM2 ectodomain to evaluate the effect of glycosylation on the thermostability of TREM2.

## RESULTS AND DISCUSSION

### Synthesis of TREM2 ectodomain via three-segment N-to-C sequential native chemical ligation

The ectodomain of TREM2(19-134) has four cysteine residues at positions 33, 51, 60, and 110 as well as two glycosylation sites at N20 and N79 (Figure 1a). We took advantage of the native cysteine residues to design a three-segment N-to-C NCL strategy with Trp50-Cys51 and Gln109-Cys110 as ligation sites (Figure 1b). Fmoc solid-phase peptide synthesis (SPPS) was used to synthesize three peptide segments ranging from 25 to 59 residues. Both TREM2(19-50)-NHNH_2_ (segment 1) and Cys-TREM2(52-109)-NHNH_2_ (segment 2) were prepared from the hydrazine resin^33, 34^, while Cys-TREM2(111-134)-NH_2_ (segment 3) was synthesized using the Rink resin. Although peptide segment 2 contains 59 residues, we were pleased to observe a major peak as the product peak in the HPLC trace of the crude peptide. The isolated yield is 2.6%. This relatively low yield is likely caused by the long polypeptide chain and the relatively high percentage (∼20%) of β-branched amino acid residues in segment 2. Another possible side reaction when synthesizing segment 2 is the C-terminal cyclization. This was previously observed when synthesizing peptide hydrazide with C-terminal Gln, although the extent of cyclization is much less compared to peptide hydrazide with C-terminal Asn.^33^ Hydrazide-based NCL^33^ was used to first ligate the segment 1 and segment 2. The resulting peptide intermediate was purified and further ligated with segment 3 to produce TREM2(19-134) with N-GlcNAc at both N20 and N79 sites. This ligation strategy gave an overall isolated yield of 28.6%. Analysis by HPLC and mass spectrometry showed high homogeneity of the final ligation product (Figures 1c and 1d).

**Figure 1.**
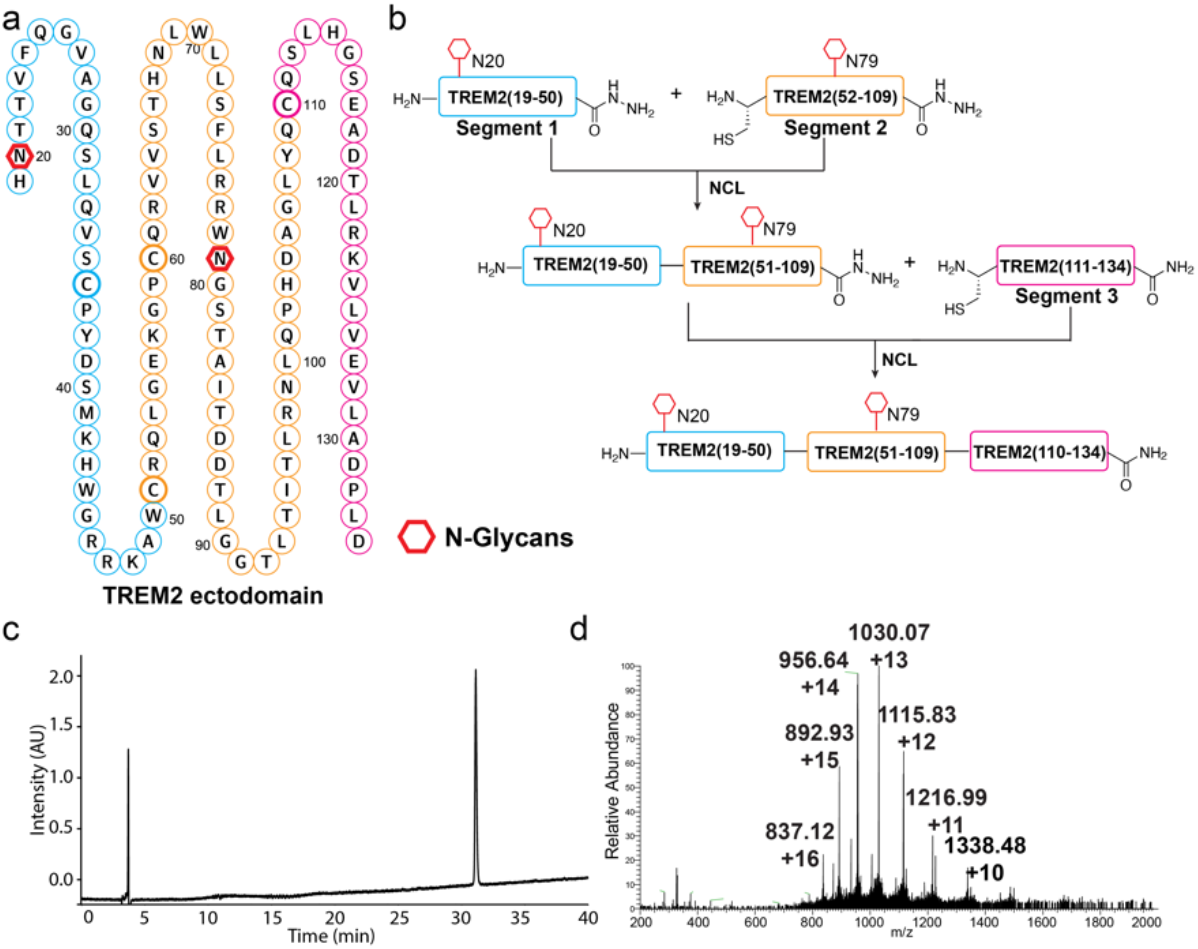
Synthesis of TREM2(19-134). (a) The sequence of TREM2 ectodomain with two glycosylation sites at N20 and N79. Three peptide fragments are color coded. (b) The synthetic scheme of three-fragment hydrazide-based native chemical ligation. Analytical HPLC trace (c) and mass spectrum (d) of synthetic TREM2(19-134) before oxidative refolding.

### Oxidative refolding of synthetic TREM2

The TREM2 ectodomain has an immunoglobulin (Ig) fold with two anti-parallel β-sheets forming a β-sandwich.^28^ The two β-sheets are bridged by a disulfide bond between C33 and C111, while another disulfide bond between C51 and C60 could provide further stability. To obtain the correctly folded protein, the synthetic TREM2(19-134) was subject to oxidative refolding and further purification by size-exclusion chromatography (Figure 2a). Analysis by analytical HPLC with a C4 column showed decreased retention time (Figure 2b), suggesting successful refolding with buried hydrophobic residues. This was also supported by mass analysis, showing a decrease of 4 Da for the refolded protein (Figure 2b). The data is consistent with the formation of two disulfide bonds. To ensure the formation of correct disulfide bonds, we also performed disulfide bond mapping. The refolded TREM2(19-134) was digested with trypsin under the non-reducing condition, and the resulting peptide mixture was analyzed by MALDI-tof mass spectrometer. We used two different MALDI matrices α-Cyano-4-hydroxycinnamic acid (CHCA) and 2,5-Dihydroxyacetophenone (DHAP). CHCA is suitable for the analysis of short peptides, while DHAP is typically used for medium-sized peptide and protein samples^35^. When using CHCA as the MALDI matrix, we observed a major peak at 1092.4, corresponding to a disulfide-bonded peptide (AW<u>C</u>R-GP<u>C</u>QR) between C51 and C60 (Figure 2c, top). With DHAP, a peak at 5452.7 was detected, consistent with another disulfide-bonded peptide between C33 and C111 (Figure 2c, bottom). No mismatched disulfide-bonded peptide was observed in this analysis. The disulfide mapping data showed the formation of correct disulfide bonds after refolding.

**Figure 2.**
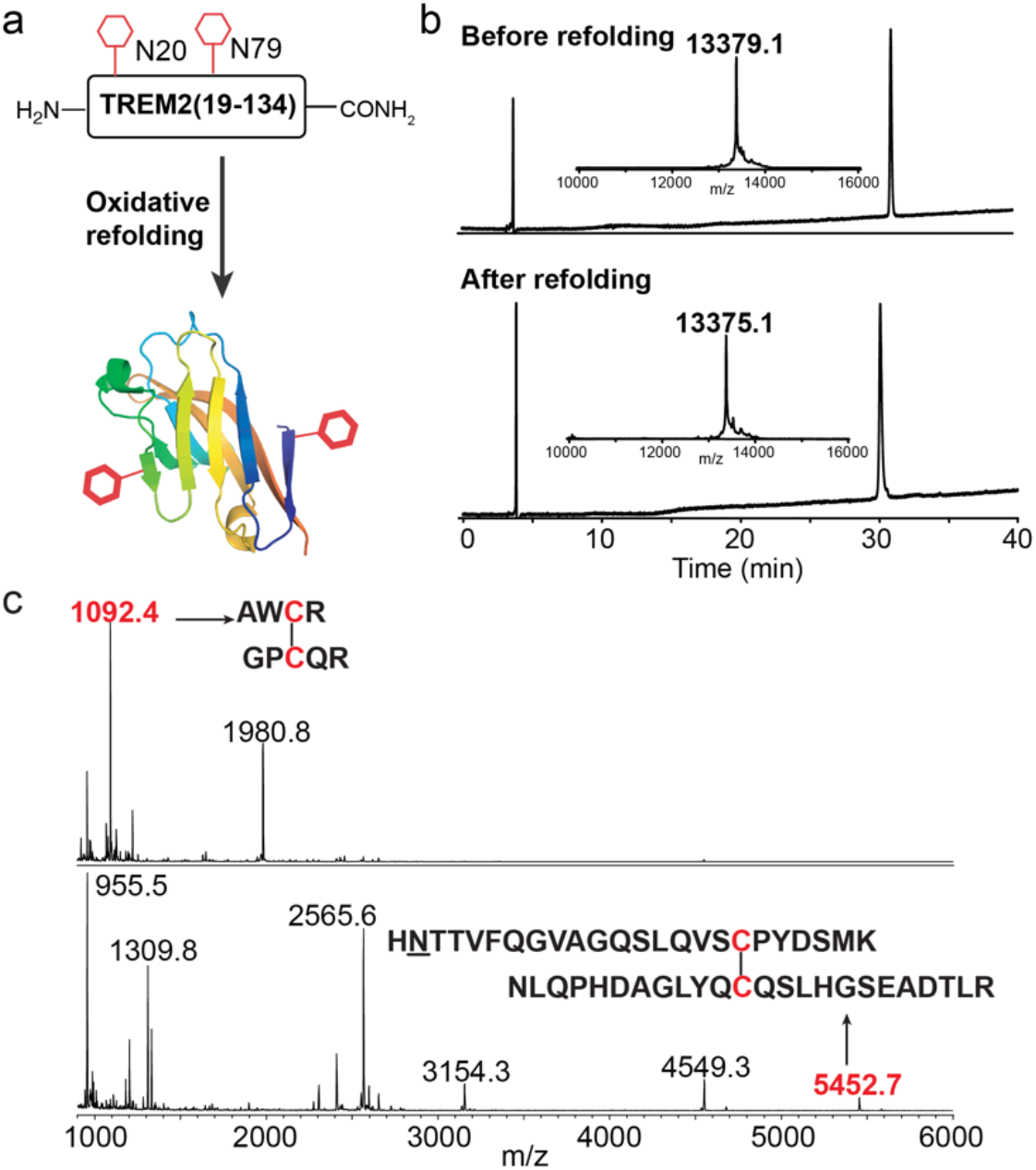
(a) Oxidative refolding of TREM2(19-134). (b) HPLC traces and mass spectra of refolded TREM2(19-134). (c) Disulfide mapping by analyzing the peptide mixture of trypsin digested TREM2(19-134). Top: CHCA as the matrix. Bottom: DHAP as the matrix.

### Structural analysis by CD and chemical cross-linking

To confirm the formation of an immunoglobulin (Ig) fold, we analyzed the protein structure by CD spectroscopy. The CD spectrum displayed double minima at 214 nm and 233 nm (Figure 3a), consistent with the reported CD spectrum of HEK293 expressed TREM2 ectodomain.^28^ Finally, we applied chemical cross-linking to determine if the refolded TREM2 can match the existing crystal structure (PDB ID 5ELI). A commercially available cross-linker BS3 was used to cross-link refolded TREM2(19-134). After reduction, alkylation and trypsin digestion, the sample was analyzed by LC-MS/MS. Plink2^36^ was used to analyze MS data to identify cross-linked peptides. The analysis yielded four cross-links that can perfectly match the crystal structures (Figure 3b). Taken together, all structural analysis supports the successful refolding of synthetic TREM2(19-134) into the Ig fold.

**Figure 3.**
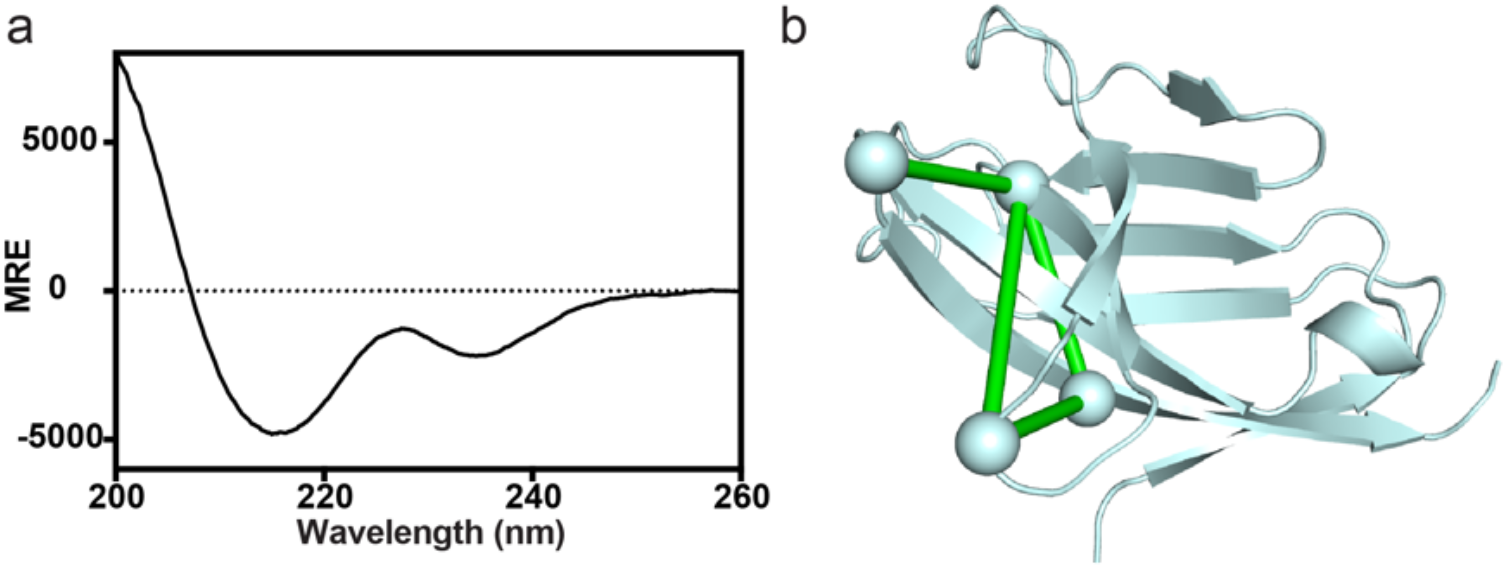
Structural characterization of refolded TREM2(19-134). (a) CD spectrum of TREM2(19-134) after refolding. (b) Cross-links mapped onto the crystal structure (PDB ID 5ELI) of TREM2(19-134).

### Refolded synthetic TREM2(19-134) enhanced microglial phagocytosis, proliferation, and survival

HMC3 is a human microglia cell line that expresses moderately low endogenous TREM2 levels and has been widely used as a model cell line to study microglial functions and the role of TREM2 in AD pathologies. Therefore, to demonstrate whether refolded synthetic TREM2(19-134) is bioactive, we treated HMC3 cells with TREM2. Under serum deprivation conditions, the addition of TREM2 enhanced the uptake of fluorescent-labeled *E. coli* bioparticles by 44.6% (Figure 4a). An Edu (5-ethynyl-2’-deoxyuridine) assay was also performed to evaluate the proliferation of microglia cells (EdU is incorporated into DNA during active DNA synthesis). After the treatment of TREM2, a 28.9% increase of microglial proliferation (Edu+ nuclei) was observed (Figure 4b). To evaluate microglia survival, cells were exposed to 100 µM H_2_O_2_ under deprivation conditions, a model of oxidative stress-induced cell death. Calcein/zombie double staining was used to distinguish viable from non-viable cells. As a result, 15% of cell death was induced by serum deprivation, and the H_2_O_2_ treatment doubled cell mortality. However, the pre-treatment of TREM2 rescued the toxicity and cell death (Figure 5). These results demonstrate that refolded synthetic TREM2(19-134) can enhance microglial phagocytosis, proliferation, and survival similar to the activities of recombinant soluble TREM2 (sTREM2) in previous studies^22, 23, 26^. These results validate the use of synthetic TREM2 constructs to investigate the functions of soluble TREM2 and the underlining molecular mechanisms.

**Figure 4.**
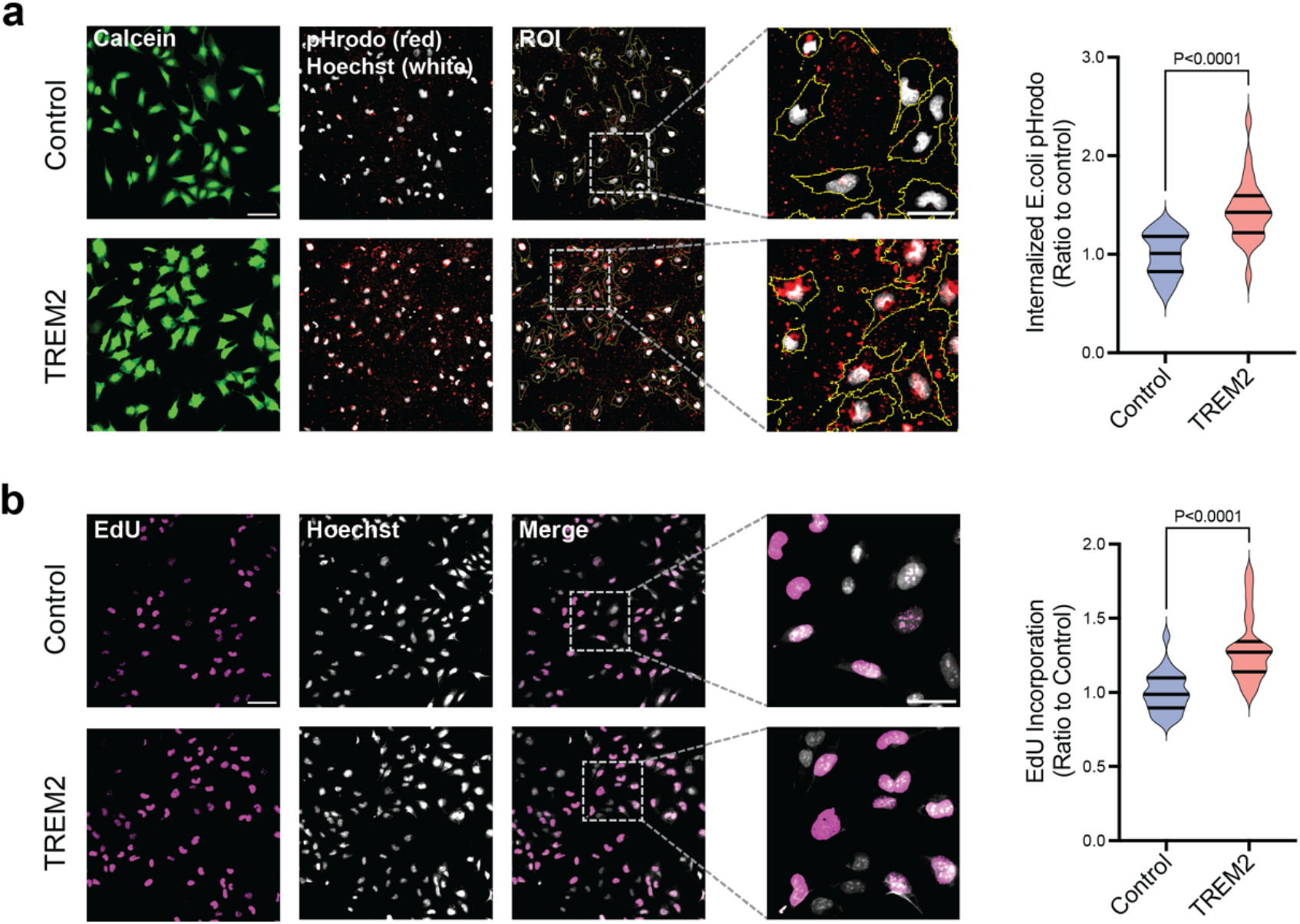
Refolded synthetic TREM2(19-134) promotes cellular proliferation and phagocytosis activities. (a) Phagocytosis assay *in vitro*. HMC3 cells were treated with 200 nM TREM2 or vehicle for 2 days under serum deprivation conditions before being exposed to pHrodo *E*.*coli* bioparticles for additional 2 hours. Representative images (left) of pHrodo (red) within the region of interest (ROI, yellow mask) delimitated by calcein (green), The total fluorescence intensity (IntDen) was measured and divided by the total number of cells. Scalebar: 100 µm for main panel, 50 µm for inset. Violin plot representation (right) of phagocytosis activity normalized to controls, showing median (middle line), 25th and 75th quartiles. P-values from unpaired t-test correspond to n = 3 combined (8 images from 2 wells, 40x magnification), statistics of each replicate in Figure S9a. (b) Proliferation assay *in vitro*. HMC3 cells were treated with 200 nM TREM2 or vehicle for 2 days under serum deprivation condition before the addition of EdU for 3 hours. Representative images (left) of EdU (magenta). Total fluorescence intensity (IntDen) was measured within the ROI delimitated by Hoechst dye (white) and divided by the total number of cells. Scalebar: 100 µm for main panel, 50 µm for inset. Violin plot representation (right) of proliferation activity normalized to controls, showing median (middle line), 25th and 75th quartiles. P-values from unpaired t-test correspond to n = 3 combined (8 images from 2 wells, 40x magnification), statistics of each replicate in Figure S9b.

**Figure 5.**
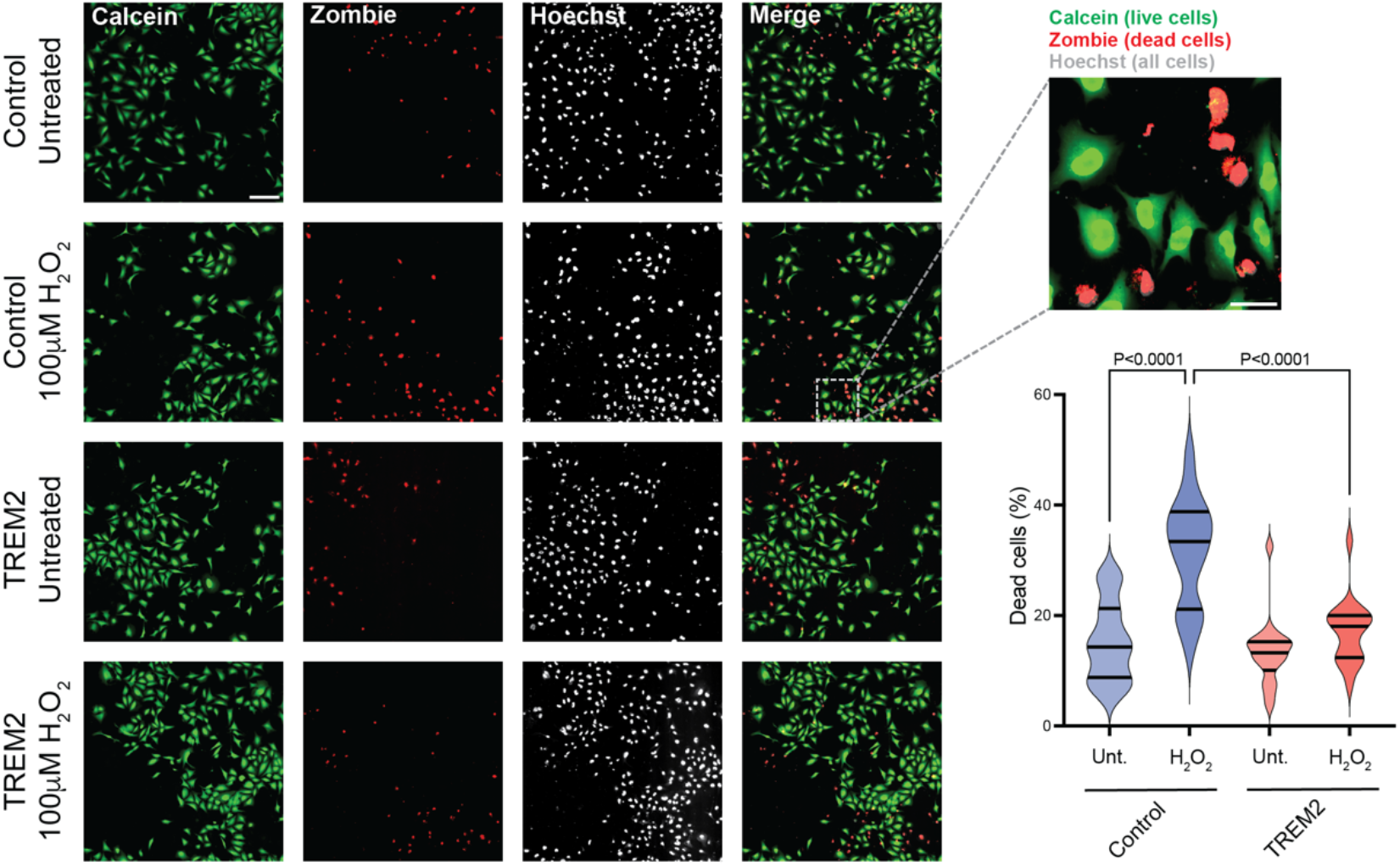
Refolded synthetic TREM2 protects against H_2_O_2_-induced cytotoxicity. HMC3 cells were treated with 200 nM TREM2 or vehicle for 1 day under serum deprivation condition before being exposed to 100 µM H_2_O_2_ for additional 24 hours. Representative images (left) of Calcein (green – live cells), Zombie dye (red – dead cells) and Hoechst (white – nuclei) are shown. Scalebar: 200 µm for main panel, 50 µm for inset. Violin plot representation (right) of the percentage of dead cells (Zombie^+^ cells per total number of cells). P-values from One-way ANOVA with Tukey post hoc test correspond to n = 3 combined (6 images from 2 wells, 20x magnification), statistics of each replicate in Figure S9c.

### Glycosylation at N79 is critical to TREM2 stability

Since glycosylation is known to enhance protein stability^37, 38^, we next tested the effect of glycosylation at N20 and/or N79 on TREM2 structure fold and thermal stability. With the established synthetic and refolding methods, we further produced two mono-glycosylated TREM2 ectodomain at either N20 (**3**) or N79 (**4**) (Figures 6a and 6b). The unglycosylated TREM2 ectodomain (**2**) was produced by *E*.*coli* expression followed by oxidative refolding^23^. Analysis by CD spectroscopy of four TREM2 ectodomain constructs revealed no significant differences (Figure 6c), suggesting that glycosylation has a minor effect on the overall TREM2 structure fold. We also measured the thermal stability of these constructs using CD spectroscopy by monitoring the mean residue ellipticity at 225 nm^28^. With elevated temperature, non-glycosylated TREM2 ectodomain (**2**) started to display a transition to the unfolded state (Figure 6d). At 50 °C, an unusual transition was observed, most likely due to aggregation. Indeed, more than half of non-glycosylated TREM2 ectodomain (**2**) aggregated after overnight incubation at 37 °C (Figure S10). By contrast, fully glycosylated TREM2 ectodomain (**1**) showed a cooperative unfolding transition with a T_m_ of 50 °C and a complete transition to the unfolded state at 62 °C (Figure 6d). The incorporation of one N-GlcNAc at N20 was able to reduce the aggregation at high temperatures. More importantly, N-GlcNAc at N79 increased the protein stability significantly. Taken together, the thermal denaturation study identified a critical role of N79 glycosylation on the thermal stability of TREM2. Interestingly, these two N glycans were shown to facilitate trafficking to the cell surface and signal transduction of full-length, membrane-tethered TREM2 in cultured cells,^39^ but the functional implications of glycosylation at N20 and N79 for soluble TREM2 ectodomain remains to be determined.

**Figure 6.**
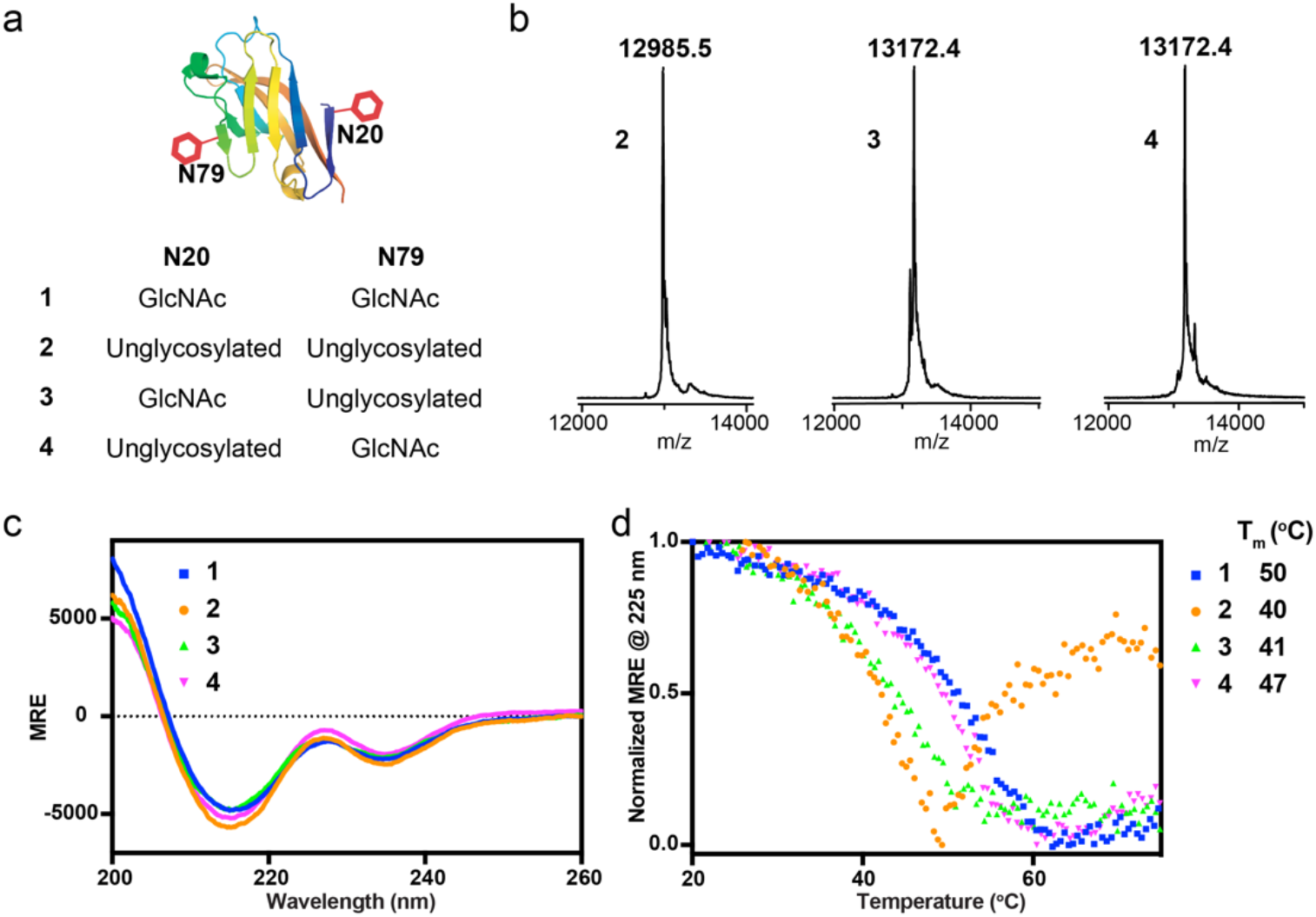
Effects of glycosylation on TREM2 structure and stability. (a) Four TREM2 ectodomain constructs with different glycosylation patterns. (b) Mass analysis of constructs **2, 3**, and **4**. (c) Comparison of CD spectra showed no significant differences. (d) Thermal stability of four constructs by CD spectroscopy.

## CONCLUSIONS

In conclusion, we have developed a method to produce synthetic TREM2 ectodomain with defined glycosylation patterns. Comprehensive structural analysis using CD spectroscopy, disulfide bond mapping, and chemical cross-linking demonstrated successful refolding of synthetic TREM2 into the Ig fold. Using human microglia HMC3 cells, we demonstrated that refolded synthetic TREM2 is bioactive and enhanced microglial phagocytosis, proliferation, and survival. With this synthetic method, we prepared TREM2 constructs with single glycosylation at either glycosylation site as well as a TREM2 construct with double glycosylation. Structural analysis using CD spectroscopy revealed that glycosylation does not affect the overall TREM2 structure fold but rather provides higher thermal stability. In particular, glycosylation at N79 is critical to TREM2 stability against aggregation. We believe this methodology will allow the introduction of not only PTMs but also other types of site-specific modifications, such as biotinylation, fluorescence labeling, affinity tags, etc. These TREM2 constructs with on-demand functions will be excellent tools to further advance our understanding of TREM2 in neurodegeneration.

## METHODS

### General information

All Fmoc protected amino acids, HCTU, DIEA, MPAA, TFA, 4-methylpiperidine, TIPS, and Fmoc-Gly-Wang resin were purchased from Chemimpex Inc. 2-chlorotrityl resin was purchased from Chempep Inc. TECP was purchased from TCI Chemicals. HPLC grade acetonitrile was purchased from VWR. All other reagents were purchased from Fisher Scientific.

### Preparation of hydrazine resin

The hydrazine resin was freshly prepared before peptide synthesis following a previously published procedure^34^. Briefly, 200 mg 2-chlorotrityl resin was swelled in 2 mL DMF for 15 min. A mixture of DIEA (125 µL) and 10% hydrazine hydrate in DMF (380 µL) was added dropwise to the resin, and the suspension was shaken at room temperature for 2 h. 2 mL MeOH was then added to the suspension, which was shaken for additional 15 min. Finally, the resin was washed with DMF (3 mL, x4).

The C-terminal amino acid was introduced to the hydrazine resin by treating resin with Fmoc-AA (4 eq.), DIEA (16 eq.) and HCTU (3.8 eq) in 2 mL DMF for 1 h at room temperature. The coupling reaction was repeated.

### Preparation of Fmoc-L-Asn((Ac)3-β-D-GlcNAc)-OH

The synthesis was done following a published procedure^40^.

### General procedure of Fmoc solid-phase peptide synthesis

#### Synthesis of peptides with C-terminal hydrazide peptides

Solid-phase peptide synthesis (SPPS) was conducted on a Purepep Chorus peptide synthesizer with the hydrazine resin (0.1 mmol scale). A typical SPPS cycle includes Fmoc deprotection (3 mL 20% 4-methylpiperidine/DMF, 10-min shaking at rt, repeated), DMF wash (4 mL, x3), coupling (4 eq. Fmoc-AA, 3.8 eq. HCTU, 8 eq. DIEA in 4 mL DMF, 120-min shaking at RT), and another DMF washing step (4 mL, x3). After SPPS cycles were completed, a final Fmoc deprotection step was conducted. The Fmoc-L-Asn((Ac)3-β-D-GlcNAc)-OH building block was used in SPPS to introduce β-GlcNAc onto N20 or/and N79 of TREM2. A final treatment of resin with 10% hydrazine in DMF (4 mL) for 16 hours at rt was done to remove acetyl groups before final peptide cleavage.

#### Synthesis of peptides with C-terminal amide

Solid-phase peptide synthesis (SPPS) was conducted on a Purepep Chorus peptide synthesizer with the Rink amide resin (0.1 mmol scale). A typical SPPS cycle includes Fmoc deprotection (3 mL 20% 4-methylpiperidine/DMF, 10-min shaking at rt, repeated), DMF wash (4 mL, x3), coupling (4 eq. Fmoc-AA, 3.8 eq. HCTU, 8 eq. DIEA in 4 mL DMF, 10-min shaking at 75 °C), and another DMF washing step (4 mL, x3). After SPPS cycles were completed, a final Fmoc deprotection step was conducted.

#### Peptide cleavage

The resin was washed with DMF (4 mL, x3) and DCM (4 mL, x5) and dried with N_2_ flow for 20 min. Peptide cleavage was done using 5 mL TFA cleavage cocktail (TFA:TIPS:H_2_O:DTT, 94:2:2:2) for 3 h. The resin slurry was filtered, and the filtrate was added dropwise to 40 mL cold diethyl ether (pre-chilled on dry ice). White precipitate was collected by centrifugation and washed with 40 mL cold ether. Crude peptide was obtained after air drying.

### General procedure of HPLC purification

The crude peptide was dissolved in 10 mL 6 M GnHCl and filtered through a 0.45 µm syringe filter. Crude material was purified by reverse-phase HPLC using a C4 preparative column (Higgins Analytical, Inc.) and solvent A (water 0.1% TFA)/B (ACN, 0.1% TFA) as the mobile phase.

### General procedure of hydrazide-based native chemical ligation

The peptide hydrazide was dissolved in an activation buffer (6 M GnHCl, 100 mM sodium phosphate, pH 3.0) to 2 mM. The solution was sonicated for 10 min and chilled on salt/ice bath for 10 min. The activation of hydrazide was done by adding freshly prepared NaNO_2_ solution (0.3 M in the activation buffer) to reach a final concentration of 30 mM. The mixture was incubated on salt/ice bath for 20 min. Then, an equal volume of a freshly prepared MPAA solution (0.2 M MPAA in 6 M GnHCl, 100 mM sodium phosphate, pH 7.5) was added to the mixture. The resulting solution was allowed to warm up to RT and added to the N-cys peptide. After 10-min sonication, the final pH was adjusted to 6.8-7.0 by addition of 2 M NaOH (increment of 2 µL). The solution was kept at RT for overnight. Upon completion, an equal volume of TCEP solution (0.1 M in GnHCl, pH 7) was added to the reaction mixture, and the solution was incubated for 10 min. The solution was acidified by mixing with TCEP solution (0.1 M in GnHCl, pH 2). After centrifugation at 14,000 g for 10 min, the supernatant was collected and purified by reverse-phase HPLC.

### Oxidative refolding and purification of TREM2

The oxidative refolding was done following a previously published procedure^29^. Briefly, after freeze drying, the synthetic TREM2(19-134) was dissolved in a denaturation buffer (8 M urea, 50 mM Tris, 10 mM DTT, pH 7.88) to 10 mg/mL. The solution was added dropwise to a pre-chilled refolding buffer (2 M urea, 50 mM Tris-HCl, 20% glycerol, 160 mM L-arginine, 3 mM cysteine, and 1 mM cystamine at pH 8.9). The solution was stirred at 4 °C for 4 hours. Upon completion (monitored by analytical HPLC), the solution was spun at 14,000 g for 40 min, and the supernatant was filtered through a 0.45 µm syringe filter. The filtrate was concentrated using a protein concentrator (3 kDa cut-off) to 2-3 mL. Further purification was done using gel filtration with 20 mM HEPES, 150 mM NaCl, 5% glycerol, pH 7.4 on an SEC column (Superdex 75 Increase 10/300 GL). Fractions containing protein were collected and stored at -80 °C.

### CD spectroscopy

Proteins (10-20 µM) were buffer exchanged to 10 mM Na phosphate, 100 mM NaF, pH 7.4 before CD analysis. Analysis was done on a Jasco 810 CD spectrometer using a 1 mm cuvette. For T_m_ experiments, 3 µM proteins were measured in a 1 cm cuvette. The ellipticity at 225 nm was monitored with increasing temperature.

### Protein stability analysis

0.4 mg/mL protein in 20 mM HEPES, 150 mM NaCl, 5% glycerol, pH 7.4 was incubated at 37 °C for overnight. After ultracentrifugation at 48 krpm using a TL55 rotor, the supernatant was removed, and the pellet was resuspended in the same volume of buffer (20 mM HEPES, 150 mM NaCl, 5% glycerol). The percentage of aggregation was determined by analyzing protein contents in the supernatant vs pellet using SDS-PAGE followed by Coomassie staining.

### Peptide/protein mass analysis

Mass data of TREM2 proteins were obtained using a Bruker autoflex MALDI-TOF mass spectrometer in a linear mode with DHAP as matrix. Mass data of peptide segments were collected on either an Agilent 6230 Tof or a Thermo Orbitrap mass spectrometer.

### Disulfide bond mapping

50 µg of refolded TREM2 protein in 100 mM ammonium bicarbonate buffer was digested with 2 µg trypsin (Sigma) at 37 °C for overnight. The resulting tryptic peptide mixture was desalted with C18 ziptip (ThermoFisher) before mass analysis using a Bruker autoflex MALDI-TOF mass spectrometer in a reflectron mode with either DHAP or CHCA as matrix.

### Chemical cross-linking

10 µM refolded TREM2 protein in the cross-linking buffer (20 mM HEPES, 150 mM NaCl, pH 7.5) was treated with 1 mM BS3 for 1 h at room temp. The reaction was quenched by adding 50 mM ammonium bicarbonate and incubating for 1 h. After quenching, protein was precipitated by mixing with six times the sample volumes of cold acetone. After incubation at -20 °C for 1 h, the suspension was spun at 13,000 *g* for 10 min to collect the pellet.

### MS analysis of cross-linked peptide

Following reduction, alkylation, and digestion of purified crosslinked protein with sequencing grade modified porcine trypsin (Promega), tryptic peptides were separated by reverse phase XSelect CSH C18 2.5 um resin (Waters) on an in-line 150 × 0.075 mm column using an UltiMate 3000 RSLCnano system (Thermo). Peptides were eluted using a 60 min gradient from 98:2 to 65:35 buffer A:B ratio. Eluted peptides were ionized by electrospray (2.4 kV) followed by mass spectrometric analysis on an Orbitrap Eclipse mass spectrometer (Thermo). MS data were acquired using the FTMS analyzer in profile mode at a resolution of 120,000 over a range of 375 to 1500 m/z. Following HCD activation, MS/MS data were acquired using the FTMS analyzer in centroid mode at a resolution of 15,000 and normal mass range with normalized collision energy of 30%. Buffer A = 0.1% formic acid, 0.5% acetonitrile. Buffer B = 0.1% formic acid, 99.9% acetonitrile. The data was analyzed using the Plink2 software^36^.

### Phagocytosis Assay

HMC3 microglia cells were plated on 384-well plate at a confluence of 1 × 10^3^ cells/well and grown in EMEM supplemented with 10% FBS and 1%Pen-Strep. On the next day, cells were washed once with serum-free medium and cultured in serum-free medium containing 200 nM of TREM2 or vehicle buffer (20 mM HEPES, 150 mM NaCl, 5% glycerol, pH 7.4) for 2 days. Cells were washed with HBSS and incubated for 2 hours with 0.1 mg/ml of pHrodo E.coli bioparticles (Invitrogen) in HBSS, cells were then washed twice and imaged immediately. 2 µM calcein and 1 μg/ml Hoechst 33342 were used to counterstain the cell bodies and nuclei, respectively. Cells were imaged with InCell Analyzer 6000 (GE Healthcare). Eight fields were captured from 2 wells, per triplicate. Fiji (ImageJ) was used to analyze the images. pHrodo fluorescence (IntDen) was measured within the Region of Interests (ROI) determined by calcein staining and divided by the total number of cells. Each replicate was normalized to their own control. Prism 9 was used for statistical analysis, unpaired t-test was used. P<0.05 was set as significant level.

### Proliferation assay

HMC3 microglia cells were cultured on 384-well plate under deprivation with 200 nM of TREM2 or vehicle buffer for 2 days. Cells were assessed using EdU Staining Proliferation Kit (Abcam) according to the manufacturer’s instructions. 1 μg/ml Hoechst 33342 was used to counterstain the nuclei. Eight fields were captured from 2 wells, per triplicate. EdU signal (IntDen) was measured within the ROI determined by Hoechst 33342 staining and divided by the total number of cells. Each replicate was normalized to their own control. Unpaired t-test was used. P<0.05 was set as significant level.

### Viability Assay

HMC3 microglia cells were cultured on 384-well plate in serum-free medium containing 200 nM of TREM2 or vehicle. H_2_O_2_ (100 µM) was added 1 day after serum deprivation and cell viability was assessed the following day by using Zombie Fixable Viability Kit (Biolegend) according to the manufacturer’s instructions. Cells were imaged immediately. 2µM calcein and 1 μg/ml Hoechst 33342 were used to counterstain the cell bodies and nuclei, respectively. Six fields were captured from 2 wells, per triplicate. Zombie positive cells were calculated and divided by the total number of cells. One-way ANOVA and Tukey post hoc tests were used. P<0.05 was set as significant level.

## ASSOCIATED CONTENT

### Supporting Information

The Supporting Information is available free of charge.

Synthesis notes; HPLC traces and mass spectra of peptide segments used in the synthesis; statistics of each biological replicate of cell-based assays; raw SDS-PAGE images of TREM2.

## AUTHOR INFORMATION

### Author Contributions

G.W., E.C., C.C., and H.W. designed research; G.W., E.C., B.H., and R.D. performed experiments; G.W., E.C., C.C., and H.W. analyzed data and wrote the paper.

## ACKNOWLEDGMENTS

The research was supported by a startup fund from Wichita State University, Ralph E. Powe Junior Faculty Enhancement Award from ORAU, and these NIH grants R15AG080493 (H.W.), RF1AG061874 (C.C.) and P01AG002132 (C.C.).

We thank Drs. Chamani Perera and Eden Go in the University of Kansas Synthetic Chemical Biology Core Facility (P20GM113117) for their help with mass analysis using MALDI-tof, Mr. Hayden Thurman for help with mass analysis using an Orbitrap mass spectrometer, and Dr. Sam Mackintosh in IDeA National Resource for Quantitative Proteomics (R24GM137786) for help with XL-MS.

## ABBREVIATIONS

AD: Alzheimer’s disease;
TREM2: Triggering receptor expressed on myeloid cells 2;
Aβ: Amyloid β;
HEK: Human Embryonic Kidney cells;
SPPS: Solid Phase Peptide Synthesis;
CD: Circular Dichroism;
NCL: Native Chemical Ligation;
N-GlcNAc: N-acetylglucosamine;
HPLC: High Performance Liquid Chromatography;
MALDI: Matrix Assisted Laser Desorption/Ionization;
CHCA: α-Cyano-4-hydroxycinnamic acid;
DHAP: 2,5-Dihydroxyacetophenone;
ECD: Ectodomain;
Ig fold: Immunoglobin fold;
AA: Amino Acid,
HCTU: *O*-(1*H*-6-Chlorobenzotriazole-1-yl)-1,1,3,3-tetramethyluronium hexafluorophosphate;
DIEA: N,N -Diisopropylethylamine;
MPAA: 4-Mercaptophenylacetic acid;
TFA: Trifluoroacetic Acid;
TIPS: Triisopropyl silane;
DMF: Dimethylformamide;
TCEP: tris(2-carboxyethyl)phosphine;
RT: Room Temperature;
GndCl: Guanidinium chloride;
DTT: Dithiothreitol;
Tris-HCl: Tris Hydrochloride;
HEPES: 2-[4-(2-Hydroxyethyl)piperazin-1-yl]ethane-1-sulfonic acid;
Edu: 5-ethynyl-2’-deoxyuridine;
ROI: region of interest;
HBSS: Hanks’ Balanced Salt Solution.

