## Supporting information for "Total Chemical Synthesis of Glycosylated TREM2 Ectodomain"

**Synthesis notes: synthesis of TREM2(19-134)**

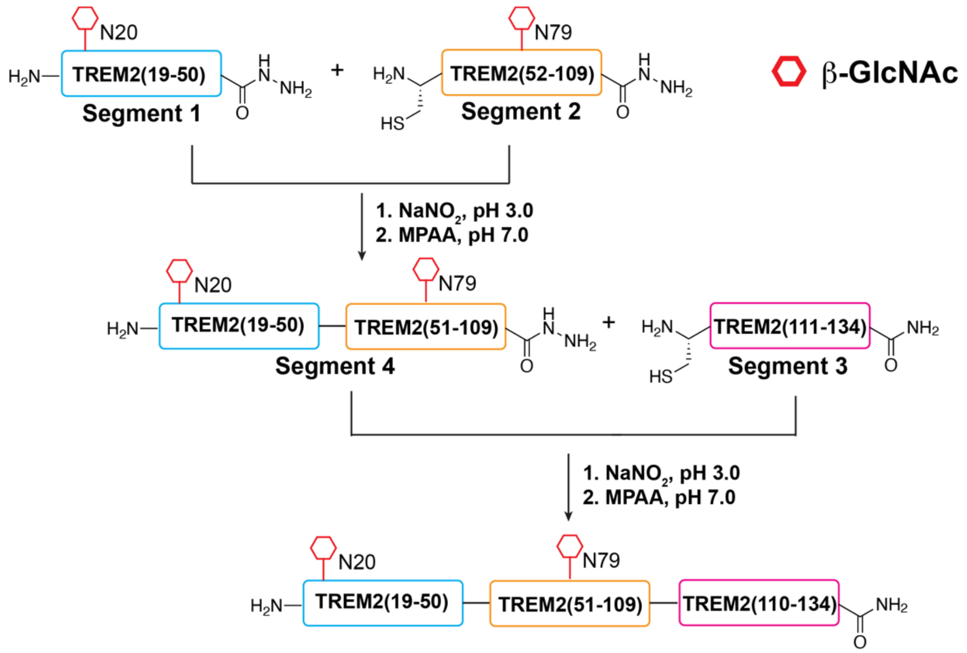


**Scheme S1.** Synthesis of TREM2(19-134).

**Synthesis of TREM2(19-50)-NHNH$_2$ (segment 1)**. The synthesis was conducted on a Purepep Chorus peptide synthesizer with the hydrazine resin (0.1 mmol scale). The C-terminal amino acid was introduced to the hydrazine resin by treating resin with Fmoc-AA (4 eq.), DIEA (16 eq.) and HCTU (3.8 eq) in 2 mL DMF for 1 h at room temperature. The coupling reaction was repeated. A typical SPPS cycle includes Fmoc deprotection (3 mL 20% 4-methylpiperidine/DMF, 10-min shaking at rt, repeated), DMF wash (4 mL, x3), coupling (4 eq. Fmoc-AA, 3.8 eq. HCTU, 8 eq. DIEA in 4 mL DMF, 120-min shaking at RT), and another DMF washing step (4 mL, x3). The coupling of Fmoc-L-Asn((Ac)3-β-D-GlcNAc)-OH was done using 2 eq. building block, 1.9 eq. HCTU, and 4 eq. DIEA in 1 mL DMF with 8-hour shaking at RT. After SPPS cycles were completed, a final Fmoc deprotection step was conducted. The acetyl group on GlcNAc was removed by treating resin with 4 mL 10% $NH_2NH_2$ in DMF for 16 h at RT. After peptide cleavage and ether precipitation, the crude peptide was purified by reverse-phase HPLC using a C4 preparative column (Higgins Analytical, Inc.) and solvent A (water 0.1% TFA)/B (ACN, 0.1% TFA) as the mobile phase at the flow rate of 13 mL/min. The peptide was separated using a linear gradient of 5% B for 2 min, 5-19.5% B in 3 min, and 19.5-34.5% B in 30 min. Yield: 22.5 mg, 5.8%.

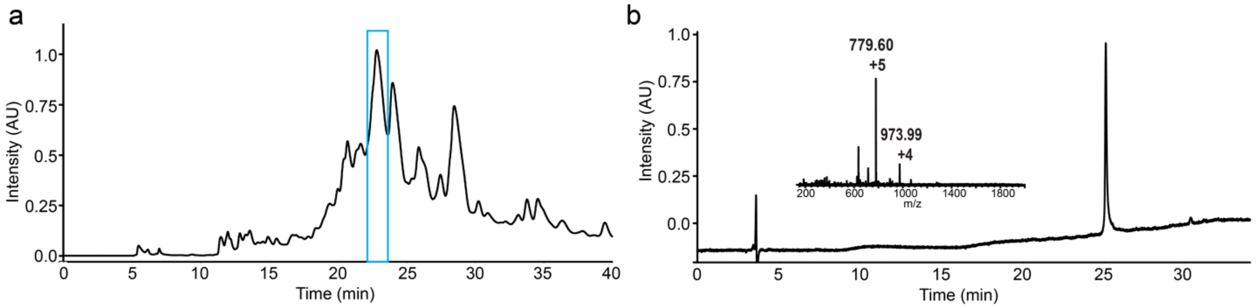


**Figure S1.** (a) Preparative HPLC trace of crude TREM2(19-50) with C-terminal hydrazide and N20-GlcNAc. Gradient: 5% B for 2 min, 5-19.5% B in 3 min, and 19.5-34.5% B in 30 min. Blue box shows the product peak. (b) Analytical HPLC trace and mass analysis of TREM2(19-50) with C-terminal hydrazide and N20-GlcNAc. Gradient: 5% B for 2 min, 5-100% B in 43 min.

**Synthesis of TREM2(51-109)-NHNH$_2$ (segment 2)**. The synthesis was conducted on a Purepep Chorus peptide synthesizer with the hydrazine resin (0.1 mmol scale). The C-terminal amino acid was introduced to the hydrazine resin by treating resin with Fmoc-AA (4 eq.), DIEA (16 eq.) and HCTU (3.8 eq) in 2 mL DMF for 1 h at room temperature. The coupling reaction was repeated. A typical SPPS cycle includes Fmoc deprotection (3 mL 20% 4-methylpiperidine/DMF, 10-min shaking at rt, repeated), DMF wash (4 mL, x3), coupling (4 eq. Fmoc-AA, 3.8 eq. HCTU, 8 eq. DIEA in 4 mL DMF, 120-min shaking at RT), and another DMF washing step (4 mL, x3). The coupling of Fmoc-L-Asn((Ac)3-β-D-GlcNAc)-OH was done using 2 eq. building block, 1.9 eq. HCTU, and 4 eq. DIEA in 1 mL DMF with 8-hour shaking at RT. After SPPS cycles were completed, a final Fmoc deprotection step was conducted. The acetyl group on GlcNAc was removed by treating resin with 4 mL 10% $NH_2NH_2$ in DMF for 16 h at RT. After peptide cleavage and ether precipitation, the crude peptide was purified by reverse-phase HPLC using a C4 preparative column (Higgins Analytical, Inc.) and solvent A (water 0.1% TFA)/B (ACN, 0.1% TFA) as the mobile phase at the flow rate of 13 mL/min. The peptide was separated using a linear gradient of 5% B for 2 min, 5-30% B in 3 min, and 30-45% B in 30 min. Yield: 17.5 mg, 2.6%.

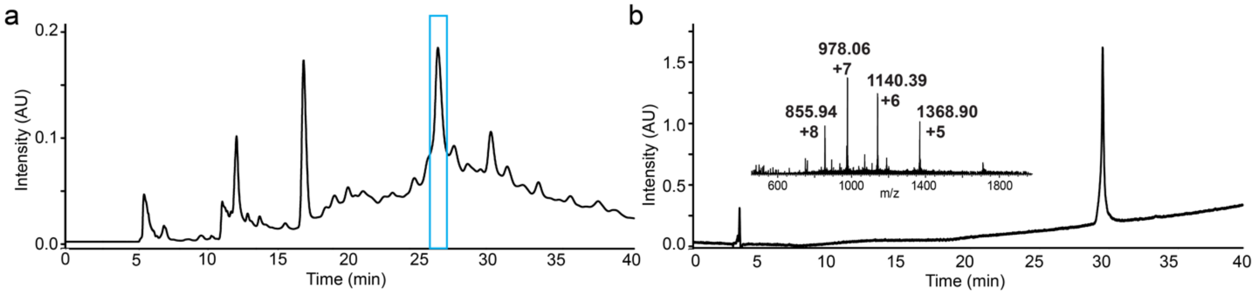


**Figure S2.** (a) Preparative HPLC trace of crude TREM2(51-109) with C-terminal hydrazide and N79-GlcNAc. Gradient: 5% B for 2 min, 5-30% B in 3 min, and 30-45% B in 30 min. Blue box shows the product peak. (b) Analytical HPLC trace and mass analysis of TREM2(51-109) with C-terminal hydrazide and N79-GlcNAc. Gradient: 5% B for 2 min, 5-100% B in 43 min.

**Synthesis of TREM2(110-134)-NH$_2$ (segment 3)**. The synthesis was conducted on a Purepep Chorus peptide synthesizer with Rink resin (0.1 mmol scale). A typical SPPS cycle includes

Fmoc deprotection (3 mL 20% 4-methylpiperidine/DMF, 10-min shaking at rt, repeated), DMF wash (4 mL, x3), coupling (4 eq. Fmoc-AA, 3.8 eq. HCTU, 8 eq. DIEA in 4 mL DMF, 120-min shaking at RT), and another DMF washing step (4 mL, x3). After SPPS cycles were completed, a final Fmoc deprotection step was conducted. After peptide cleavage and ether precipitation, the crude peptide was purified by reverse-phase HPLC using a C4 preparative column (Higgins Analytical, Inc.) and solvent A (water 0.1% TFA)/B (ACN, 0.1% TFA) as the mobile phase at the flow rate of 13 mL/min. The peptide was separated using a linear gradient of 5% B for 2 min, 5-30% B in 3 min, and 30-45% B in 30 min. Yield: 32 mg, 12%.

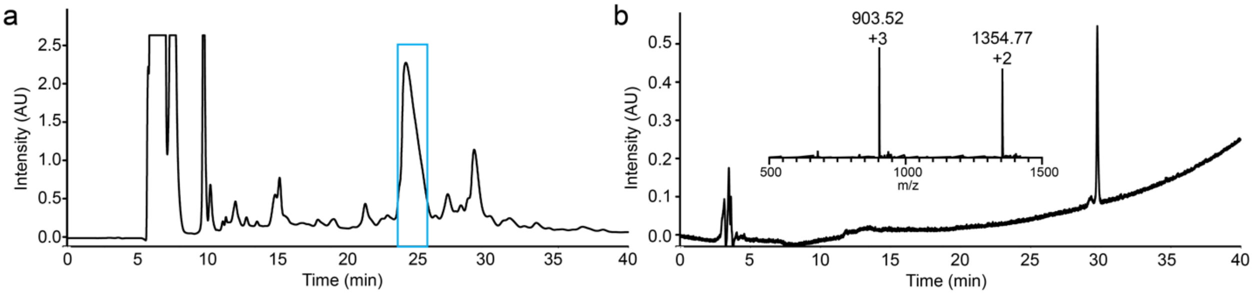


**Figure S3.** (a) Preparative HPLC trace of crude TREM2(110-134) with C-terminal amide. Gradient: 5% B for 2 min, 5-30% B in 3 min, and 30-45% B in 30 min. Blue box shows the product peak. (b) Analytical HPLC trace and mass analysis of TREM2(110-134) with C-terminal amide. Gradient: 5% B for 2 min, 5-100% B in 43 min.

**Ligation of segment 1 and segment 2 to prepare segment 4.** Segment **1** (15 mg) was dissolved in 1.2 mL activation buffer (6 M GnHCl, 100 mM sodium phosphate, pH 3.0). The solution was sonicated for 10 min and chilled on salt/ice bath for 10 min. The activation of hydrazide was done by adding 120 μL freshly prepared $NaNO_2$ solution (0.3 M in the activation buffer) to reach a final concentration of 30 mM. The mixture was incubated on salt/ice bath for 20 min. Then, 1.2 mL freshly prepared MPAA solution (0.2 M MPAA in 6 M GnHCl, 100 mM sodium phosphate, pH 7.5) was added to the mixture. The resulting solution was allowed to warm up to RT and added to segment **2** (25 mg). After 10-min sonication, the final pH was adjusted to 6.8-7.0 by addition of 4 M NaOH (increment of 2 μL). The solution was kept at RT for 12 h. Upon completion, 3 mL TCEP solution (0.1 M in GnHCl, pH 7) was added to the reaction mixture, and the solution was incubated for 10 min. The solution was acidified by mixing with 40 mg TCEP. After filtration with a syringe filter, the filtrate was collected and purified by reverse-phase HPLC using a C4 preparative column (Higgins Analytical, Inc.) and solvent A (water 0.1% TFA)/B (ACN, 0.1% TFA) as the mobile phase at the flow rate of 13 mL/min. The peptide was separated using a linear gradient of 5% B for 2 min, 5-27% B in 3 min, and 27-42% B in 30 min. Yield: 22 mg, 55%.

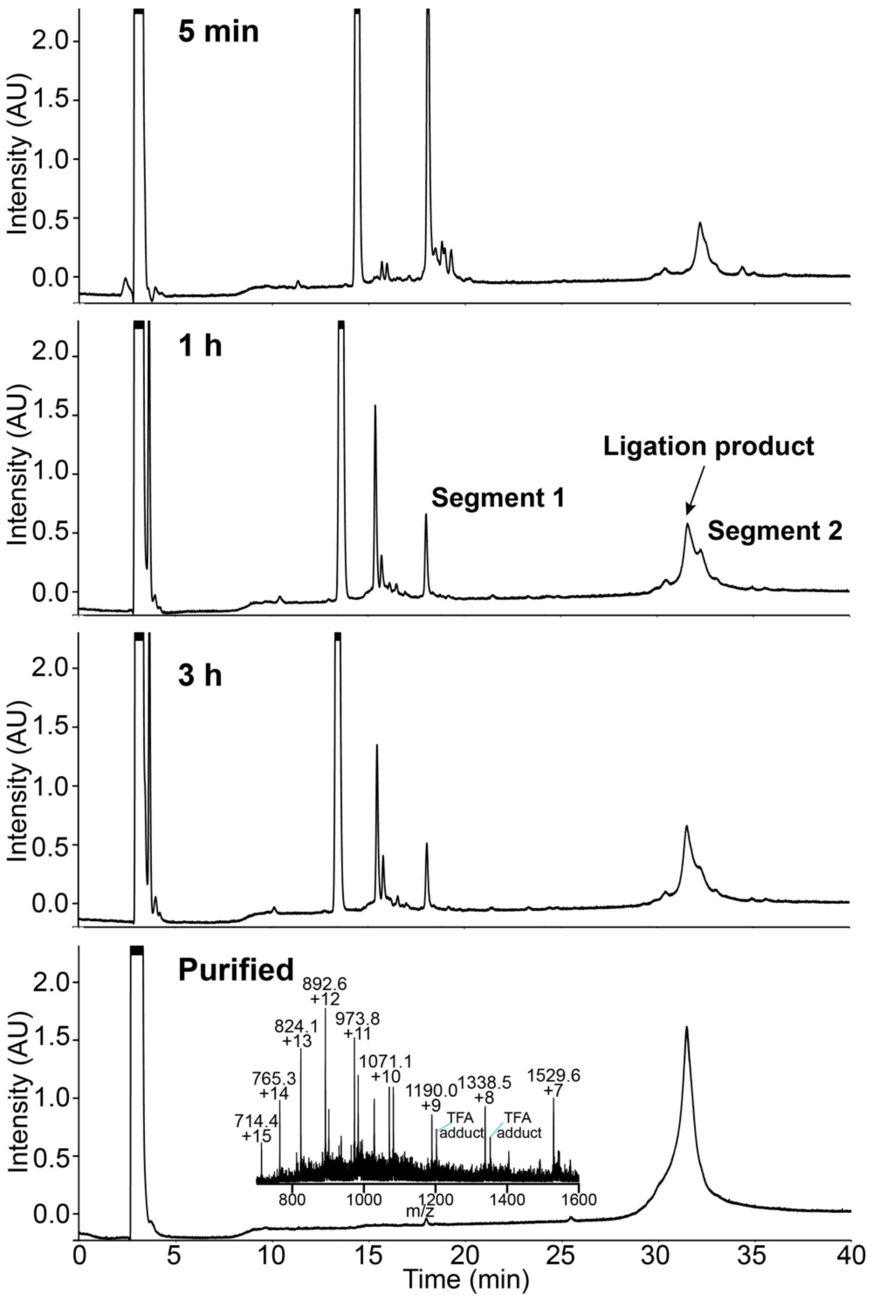


**Figure S4.** Ligation of segment **1** and segment **2** at different time points. Gradient: 5% B for 2 min, 5-30% B in 3 min, 30-45% B in 30 min.

**Ligation of segment 3 and segment 4.** Segment **4** (10 mg) was dissolved in 0.4 mL activation buffer (6 M GnHCl, 100 mM sodium phosphate, pH 3.0). The solution was sonicated for 10 min and chilled on salt/ice bath for 10 min. The activation of hydrazide was done by adding 40 μL freshly prepared $NaNO_2$ solution (0.3 M in the activation buffer) to reach a final concentration of 30 mM. The mixture was incubated on salt/ice bath for 20 min. Then, 0.4 mL freshly prepared MPAA solution (0.2 M MPAA in 6 M GnHCl, 100 mM sodium phosphate, pH 7.5) was added to the mixture. The resulting solution was allowed to warm up to RT and added to segment **3** (3.5 mg). After 10-min sonication, the final pH was adjusted to 6.8-7.0 by addition of 1 M NaOH (increment of 2 μL). The solution was kept at RT for 9 h. Upon completion, 0.8 mL TCEP solution (0.1 M in GnHCl, pH 7) was added to the reaction mixture, and the solution was incubated for 10 min. The solution was acidified by mixing with 20 mg TCEP. After filtration with a syringe filter, the filtrate was collected and purified by reverse-phase HPLC using a C4 preparative column (Higgins Analytical, Inc.) and solvent A (water 0.1% TFA)/B (ACN, 0.1% TFA) as the mobile phase at the flow rate of 13 mL/min. The peptide was separated using a linear gradient of 5% B for 2 min, 5-27% B in 3 min, and 27-42% B in 30 min. Yield: 6.5 mg, 52%.

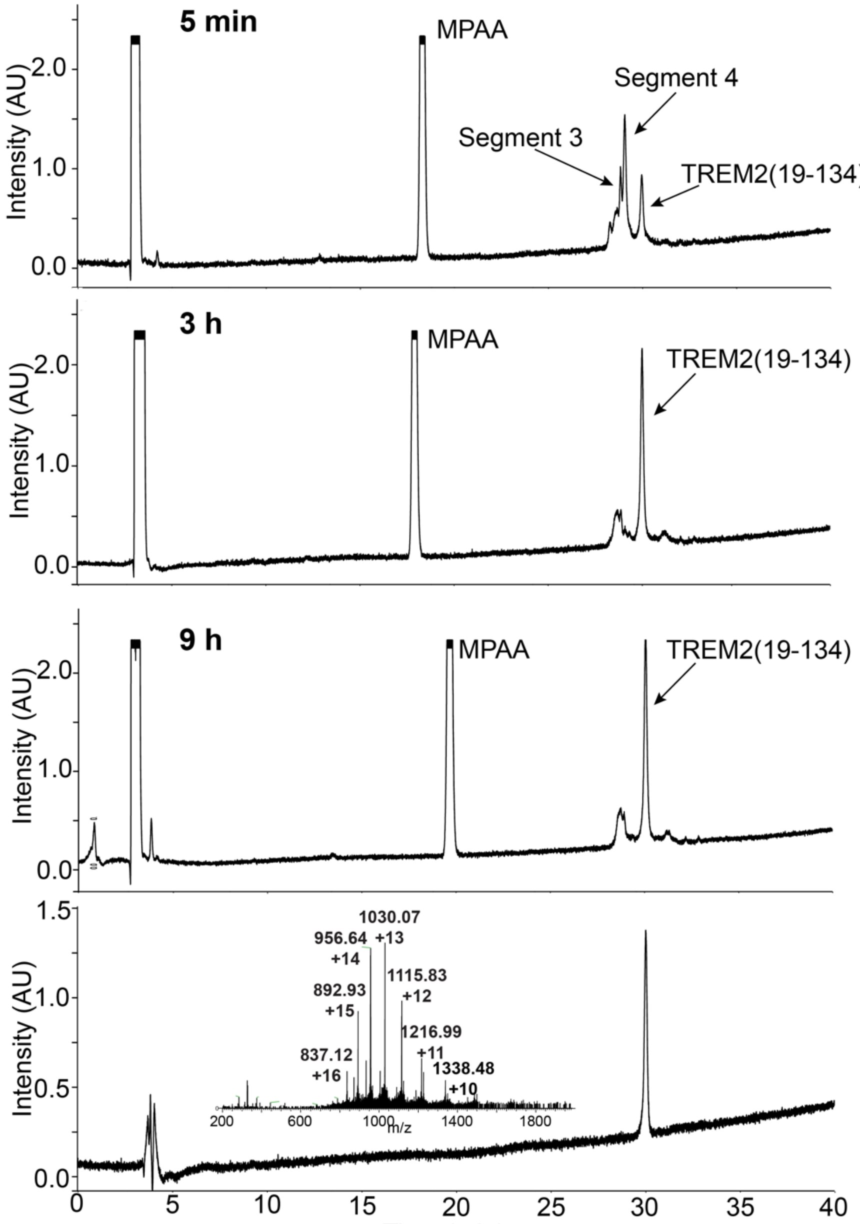


**Figure S5.** Ligation of segment **3** and segment **4** at different time points. Gradient: 5% B for 2 min, 5-100% B in 43 min.

**Refolding of synthetic TREM2(19-134)**. After freeze drying, 6.5 mg synthetic TREM2(19-134) was dissolved in 2.6 mL denaturation buffer (8 M urea, 50 mM Tris, 10 mM DTT, pH 7.88) to 2.5 mg/mL. The solution was added dropwise to 62.4 mL pre-chilled refolding buffer (2 M urea, 50 mM Tris-HCl, 20% glycerol, 160 mM L-arginine, 3 mM cysteine, and 1 mM cystamine at pH 8.9). The solution was stirred at 4 °C. Upon completion (monitored by analytical HPLC), the solution was spun at 14,000 g for 40 min, and the supernatant was filtered through a 0.45 µm syringe filter. The filtrate was concentrated using a protein concentrator (3 kDa cut-off) to 2-3 mL. Further purification was done using gel filtration with 20 mM HEPES, 150 mM NaCl, 5% glycerol, pH 7.4 on an SEC column (Superdex 75 Increase 10/300 GL). Fractions containing monomeric protein was collected, concentrated to 53.5 µM (1.5 mL final volume) and stored at -80 °C. Yield: 18.8%.

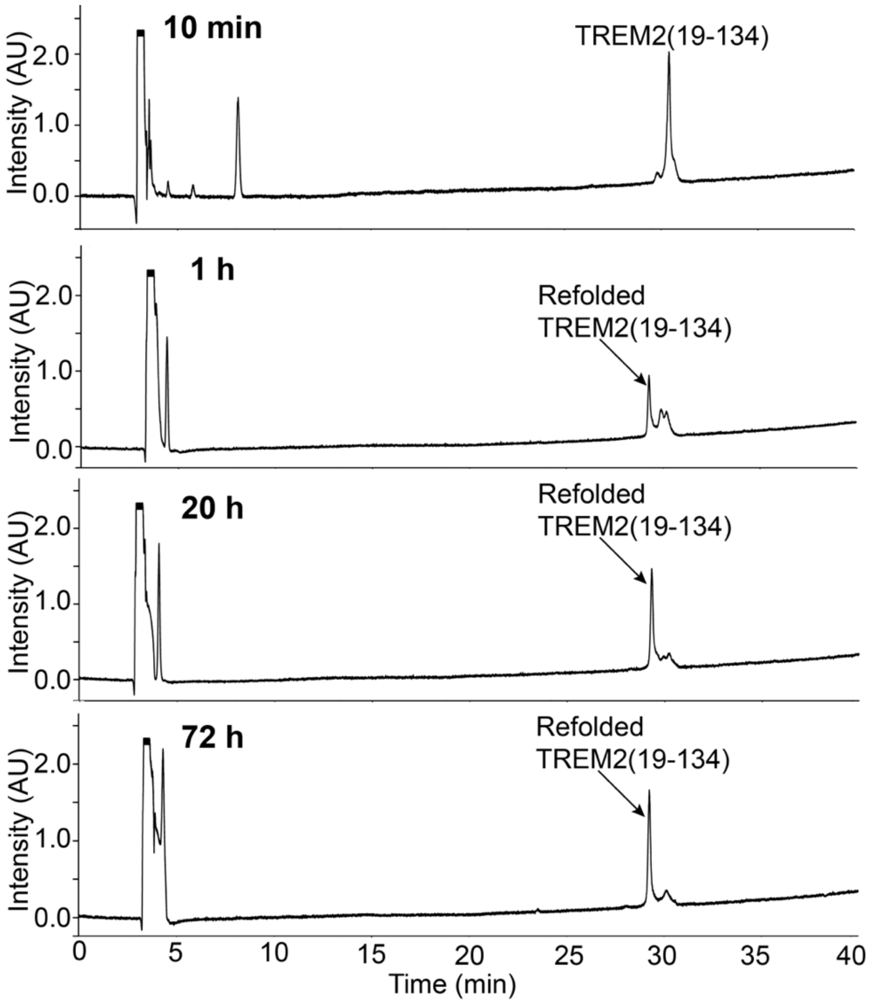


**Figure S6.** Refolding of synthetic TREM2(19-134) at different time points. Gradient: 5% B for 2 min, 5-100% B in 43 min.

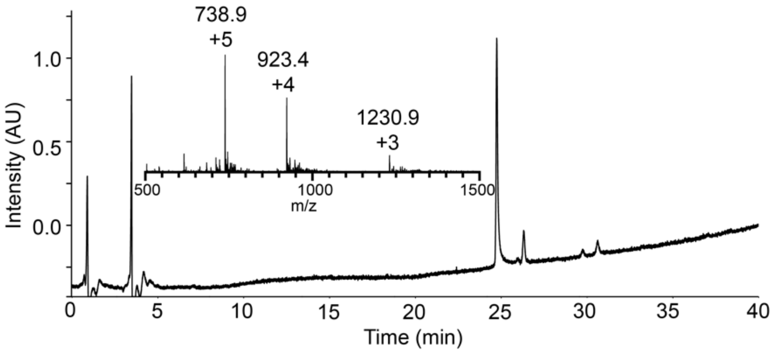


**Figure S7.** Analytical HPLC trace and mass analysis of TREM2(19-50) with C-terminal hydrazide.

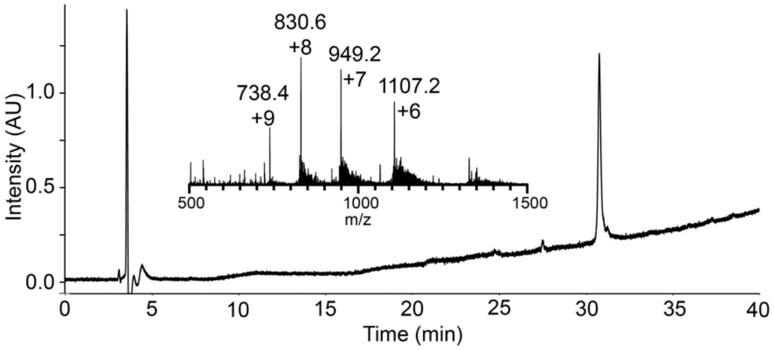


**Figure S8**. Analytical HPLC trace and mass analysis of TREM2(51-109) with C-terminal hydrazide.

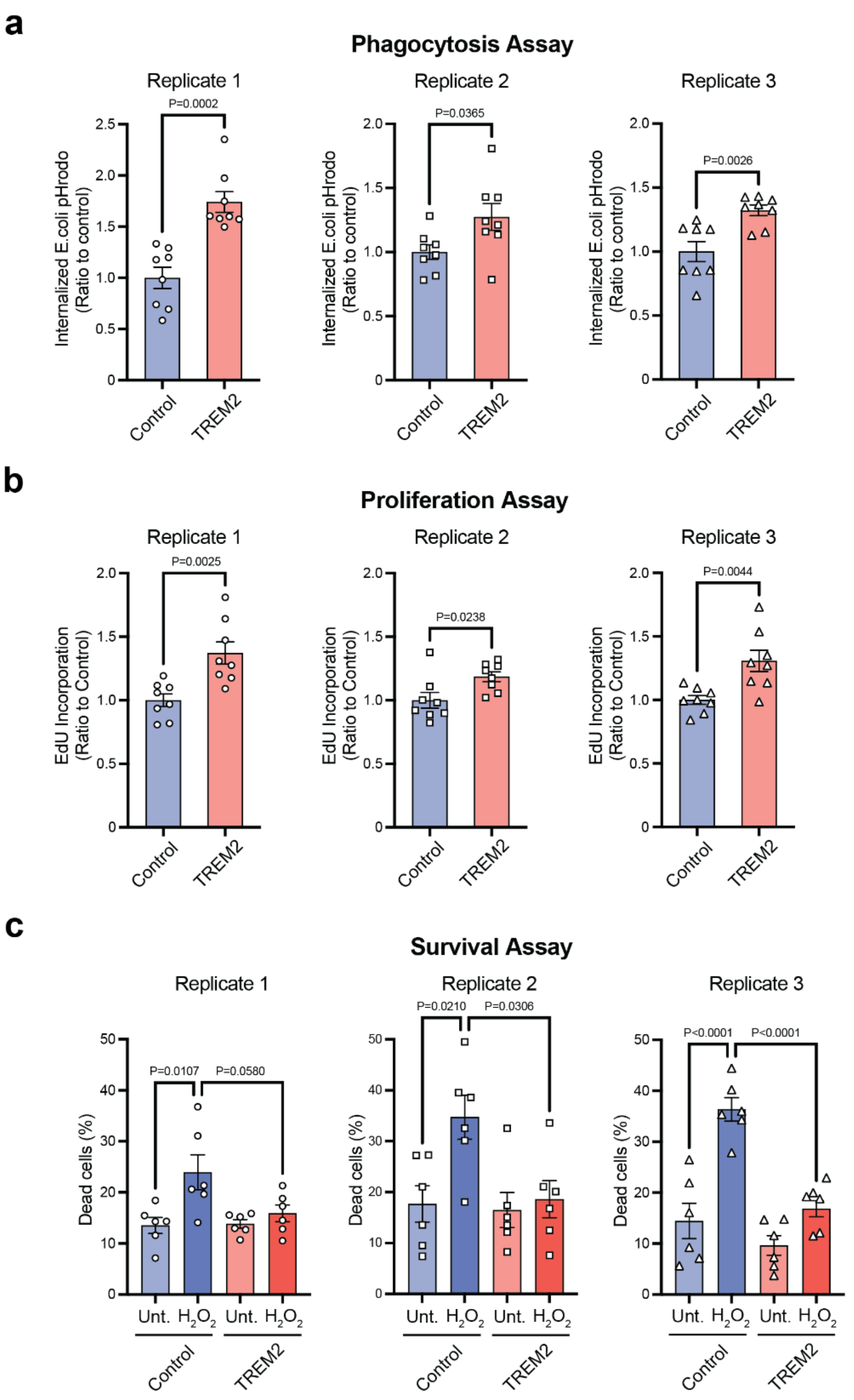


**Figure S9. Independent biological replicate experiments for all cellular assays.** Quantification of phagocytosis (a), proliferation (b) and survival (c) assays expressed as mean ± SEM. Each symbol dot indicates a technical replicate. Unpaired t-test for phagocytosis and proliferation assays. One-way ANOVA with Tukey post hoc for survival assay.

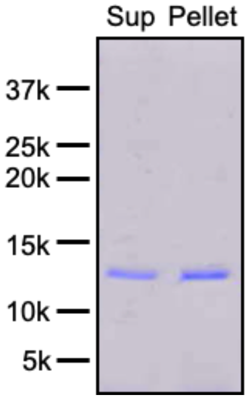


**Figure S10.** Aggregation of non-glycosylated TREM2(19-133) after overnight incubation at 37 °C.
